# DLDG1, the Arabidopsis Homolog of the Cyanobacterial H^+^-Extrusion Protein, Regulates Non-Photochemical Quenching in Chloroplasts

**DOI:** 10.1101/731653

**Authors:** Kyohei Harada, Takatoshi Arizono, Ryoichi Sato, Mai Duy Luu Trinh, Akira Hashimoto, Masaru Kono, Masaru Tsujii, Nobuyuki Uozumi, Shinichi Takaichi, Shinji Masuda

## Abstract

Plants convert solar energy into chemical energy through photosynthesis, which supports almost all life activities on earth. Because the intensity and quality of sunlight can change dramatically throughout the day, various regulatory mechanisms help plants adjust their photosynthetic output accordingly, including the regulation of light energy accumulation to prevent the generation of damaging reactive oxygen species. Non-photochemical quenching (NPQ) is a regulatory mechanism that dissipates excess light energy, but how it is regulated is not fully elucidated. Herein, we report a new NPQ-regulatory protein named Day-Length-dependent Delayed-Greening1 (DLDG1). The Arabidopsis DLDG1 associates with the chloroplast envelope membrane, and the *dldg1* mutant had a large NPQ value compared with wild type. The mutant also had a pale-green phenotype in developing leaves but only under continuous light; this phenotype was not observed when *dldg1* was cultured in the dark for ≥8 h per day. DLDG1 is a homolog of the plasma-membrane-localizing cyanobacterial proton-extrusion-protein A that is required for light-induced H^+^ extrusion, and also shows similarity in its amino-acid sequence to that of Ycf10 encoded in the plastid genome. Arabidopsis DLDG1 enhances the growth-retardation phenotype of the *Escherichia coli* K^+^/H^+^ antiporter mutant, and the everted membrane vesicles of the *E. coli* expressing DLDG1 show the K^+^/H^+^ antiport activity. Our findings suggest that DLDG1 functionally interacts with Ycf10 to control H^+^ homeostasis in chloroplasts, which is important for the light-acclimation response, by optimizing the extent of NPQ.

## Introduction

Plants convert light energy into chemical energy by photosynthesis. Because sunlight is the only energy source naturally supplied to the earth in large quantities, photosynthesis supports usable energy for almost all life activities on earth. Photosynthesis occurs via two major steps: an electron-transfer reaction and a carbon-fixing reaction. The electron-transfer reaction, which extracts electrons from water, is catalyzed by photosystem II (PSII). The electrons are then transferred from plastoquinone, to the cytochrome *b*_*6*_*f* complex, to plastocyanin, to photosystem I (PSI), and finally to ferredoxin on thylakoid membranes. The light-harvesting chlorophyll-protein complexes (LHCs) I and II absorb light energy and transfer most of the captured energy to PSI and PSII, respectively. Reduced ferredoxin potentiates the activity of the ferredoxin-NADP^+^ oxidoreductase to reduce NADP^+^ to NADPH. In this process, a proton motive force across the thylakoid membrane (*pmf*) is generated that is composed of a H^+^-concentration gradient (ΔpH) and an electric potential (**ΔΨ**). In turn, ΔpH and ΔΨ contribute equally to ATP synthesis by plastidial ATP synthase embedded in the thylakoid membrane. The NADPH and ATP thus generated are used for CO_2_ fixation by the carbon-fixing reaction (Calvin-Benson cycle) (Blankenship, 2002).

Under natural conditions, the intensity and quality of sunlight will change dramatically over a 24-h period and whether clouds are present or absent. To survive under such variable environments, plants have acquired sophisticated mechanisms to regulate photosynthesis in response to different light conditions. Plants have mechanisms that dissipate excessive light energy as heat or chemical energy to acclimate to intense light. One such mechanism is non-photochemical quenching (NPQ) (Ruban, 2016). When this mechanism is missing, excess light energy induces the generation of reactive oxygen species, which can damage plant cells (Niyogi, 1999; Krieger-Liszkay, 2004; Møller et al., 2007). In particular, the aforementioned complexes involved in the electron-transfer reaction are susceptible to damage by the reactive oxygen species produced under intense light (Aro et al., 1993; Nishiyama et al., 2001; Tiwari et al., 2016). Therefore, NPQ, under intense light, must be precisely controlled to prevent cell damage while maintaining photosynthetic efficiency.

NPQ reactions can occur over different relaxation-time scales depending on the molecular players involved and include pH-regulated energy-dissipation in the LHCII (qE) (Krause et al., 1982), state transitions (qT) (Dietzel et al., 2008), chloroplast movement– dependent quenching (qM) (Cazzaniga et al., 2013), photoinhibition of PSII (qI) (Krause, 1988), and lipocalin-dependent quenching (qH) (Malnoë et al., 2018). qE is the major component of NPQ in land plants and is regulated by xanthophyll-cycle carotenoids whose composition is controlled by extent of ΔpH. Upon transition from weak to intense light, violaxanthin is converted into zeaxanthin by violaxanthin de-epoxidase (VDE) in the chloroplast. Violaxanthin absorbs light energy and transfers that energy to chlorophyll (Chl), whereas zeaxanthin receives energy from excited Chl molecules and dissipates it as heat (Siefermann-Harms, 1987; Demmig-Adams, 1990) through leading LHCII aggregation (Johnson et al., 2011); the zeaxanthin-dependent NPQ is also referred to as qZ (Nilkens et al., 2010). Therefore, reversible conversion of xanthophyll according to the environmental light intensity is important to prevent cell damage caused by excessive light energy and to maximize photosynthesis under weak light. VDE is exposed to the lumen, and its enzymatic activity is enhanced by protonation of certain amino acids that is accelerated by formation of the ΔpH through photosynthetic electron transfer (Hieber et al., 2002; Jahns et al., 2009). A mutation in *NPQ4*, which encodes the thylakoid membrane protein PsbS, results in an abnormal decrease in qE (Li et al., 2000), indicating that PsbS is necessary for qE. A recent study indicated that PsbS interacts with specific LHCII complexes for qE induction, which is affected by the zeaxanthin level and the magnitude of ΔpH across the thylakoid membrane (Sacharz et al., 2017).

Although the mechanisms of qE induction are well understood, the detailed mechanisms by which the lumen pH is appropriately adjusted according to light intensity have yet to be elucidated. We recently screened for Arabidopsis genes that are co-expressed with NPQ-related genes that are specifically conserved in oxygenic phototrophs; we identified a gene that encodes an uncharacterized NPQ-regulatory protein, which we named *Fluctuating-Light Acclimation Protein1* (*FLAP1*) (Sato et al., 2017). The Arabidopsis *flap1* mutant has a larger NPQ value than does the wild type (WT) and has a pale-green leaf phenotype, but only when grown under light of fluctuating intensity (fluctuating light) (Sato et al., 2017; Trinh et al., 2019), indicating that FLAP1 helps plants acclimate to fluctuating light. For the study reported herein, with the use of reverse genetic screening, we identified another open-reading frame, namely *Day-Length-dependent Delayed-Greening1* (*DLDG1*), as a new NPQ-regulatory component specifically conserved in oxygenic phototrophs. Notably, the DLDG1 homolog proton-extrusion-protein A (PxcA) from the cyanobacterium *Synechocystis* sp. PCC6803 has been shown to be involved in light-induced H^+^ extrusion (Katoh et al., 1996; Sonoda et al., 1998). We propose that plant DLDG1 homologs control H^+^ homeostasis in chloroplasts for acclimation to changes in light intensity.

## Results

### Identification of a candidate gene responsible for controlling NPQ

We screened for Arabidopsis genes that have the following properties: (i) predicted to be chloroplastic, (ii) co-expressed with known NPQ-related genes, and (iii) conserved in oxygenic phototrophs. In addition to FLAP1, the details of which we reported previously (Sato et al., 2017; Trinh et al., 2019), we identified, by reverse genetic screening, another candidate NPQ-related gene at the open-reading frame, At4g31040, which we abbreviated as *DLDG1*. Examination of the co-expression database ATTEDII (Obayashi et al., 2018) revealed that *DLDG1* is co-expressed with *NPQ1* (encodes VDE), *FTRA2* (ferredoxin-thioredoxin oxidoreductase subunit A), and *FNR2* (ferredoxin-NADP^+^ oxidoreductase 2), whose functions seem to be important for controlling NPQ and ΔpH. DLDG1 consists of 438 amino acid residues and contains an N-terminal chloroplast-type transit peptide and three transmembrane regions, as revealed by ChloroP (http://www.cbs.dtu.dk/services/ChloroP) and SOSUI (http://harrier.nagahama-i-bio.ac.jp/sosui/sosui_submit.html), respectively (Fig. 1A). A search for similar sequences in the genomes of other oxygenic phototrophs in different phyla, including cyanobacteria, identified DLDG1 homologs. Two different types of DLDG1 homologs were found in these genomes, with one being encoded in the plastid genome of plant cells and annotated as *Ycf10* homologs. An amino-acid sequence alignment indicated that Ycf10-type proteins contain a C-terminal region similar in sequence to that of DLDG1 (Supplementary Fig. S1), which is a member of the CemA superfamily in the domain database InterPro (https://www.ebi.ac.uk/interpro/) (Fig. 1A). Phylogenetic analysis revealed that plant DLDG1s and Ycf10s are clearly separated from each other evolutionarily (based on bootstrap values) (Supplementary Fig. S2). One of the two DLDG1/Ycf10 homologs in the cyanobacterium *Synechocystis* sp. PCC6803 (PxcA; slr1596) was shown to be localized in the plasma membrane and shown to be essential for H^+^ extrusion into the medium upon illumination with light (Katoh et al., 1996; Sonoda et al., 1998), suggesting that plant-type DLDG1/Ycf10 homologs might be involved in controlling chloroplast H^+^ homeostasis.

**Fig. 1.**
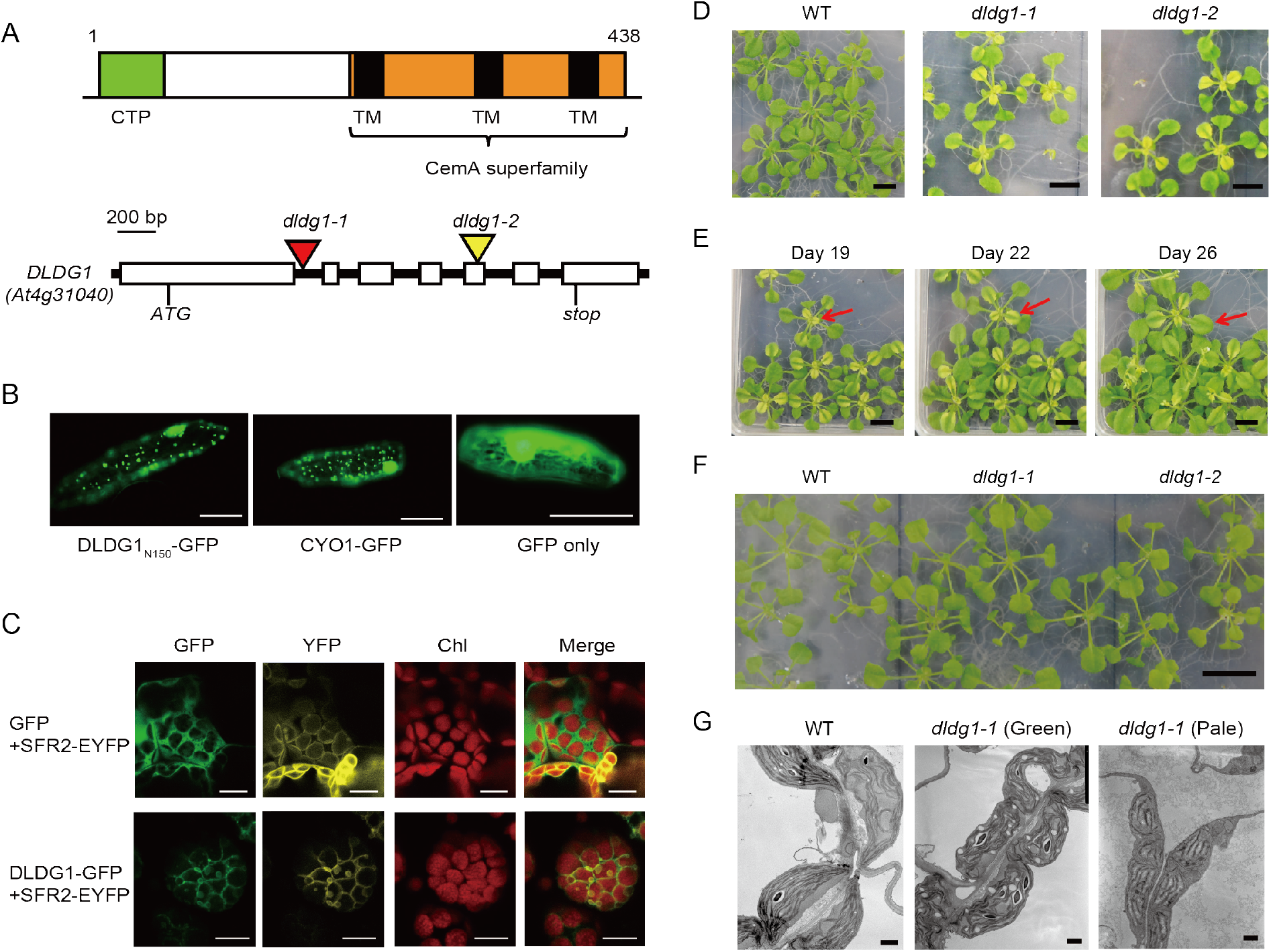
Characterization of DLDG1. (A) Schematic representation of DLDG1 and its gene structure. (Top) The predicted chloroplast transit peptide (CTP) and transmembrane regions (TM) in DLDG1 are indicated. (Bottom) The positions of the two mutant alleles, the initiation codon (ATG), and the stop codon in *DLDG1* are depicted. (B) Localization of GFP fused to the N-terminal 150 residues of DLDG1 (DLDG1_N150_-GFP). The plastid-localized CYO1-GFP (Shimada et al., 2007) and GFP served as the positive and negative controls, respectively. Scale bar, 100 μm. (C) Localization of DLDG1-GFP or GFP co-expressed with SFR2-EYFP that localizes in the chloroplast envelope membrane(Fourrier et al., 2008). Scale bar, 10 μm. (D) Phenotypes of leaves from 19-day-old WT and *dldg1* plants grown under continuous light (50 μmol m^−2^ s^−1^). Scale bar, 1 cm. (E) Phenotypes of leaves from 19-, 22-, and 26-day-old plants and a *dldg1* plant (*dldg1-2*) grown under continuous light (50 μmol m^−2^ s^−1^). Red arrows indicate the same leaf at different stages. Scale bar, 1 cm. (F) Phenotypes of leaves from 19-day-old WT and *dldg1* plants grown under a 12-hr dark/12-hr light (50 μmol m^−2^ s^−1^) cycle. Scale bar, 1 cm. (G) Ultrastructures of chloroplasts in WT leaves and in mature green leaves and young pale-green leaves from a *dldg1-1* plant grown under continuous light (50 μmol m^−2^ s^−1^). Scale bar, 1 μm.

### DLDG1 localizes to the chloroplast envelope membrane in Arabidopsis

To characterize the transit peptide of DLDG1, the recombinant gene encoding green fluorescent protein (GFP) attached to the N-terminal 150 residues of DLDG1 (DLDG1_N150_-GFP) was introduced into onion epidermal cells via a particle gun to transiently express the protein. We also expressed CYO1-GFP and GFP as positive and negative controls, respectively, for plastid localization (Shimada et al., 2007). As shown in Fig. 1B, the DLDG1_N150_-GFP fluorescence signal was similar to that of CYO1-GFP but not that of GFP, which indicated that DLDG1 has a chloroplast transit peptide at its N-terminus.

A construct containing GFP fused to full-length DLDG1 (DLDG1-GFP) was co-expressed in tobacco leaves with SFR2-EYFP, which localizes to the chloroplast envelope membrane (Fourrier et al., 2008). GFP was also co-expressed with SFR2-EYFP as a negative control. As shown in Fig. 1C, Chl autofluorescence was surrounded by DLDG1-related GFP signals, which clearly overlapped with those of SFR2-EYFP. Conversely, isolated GFP signals were seen in the cytoplasm and did not overlap with those of SFR2-EYFP, suggesting that DLDG1 localizes to the chloroplast envelope membrane.

### Phenotypes of Arabidopsis *dldg1* plants

The T-DNA insertional Arabidopsis DLDG1 mutants (*dldg1-1* and *dldg1-2*) were obtained from the Arabidopsis Biological Resource Center. PCR-based genotyping indicated that *dldg1-1* and *dldg1-2* have T-DNA insertions in the intron 1 and the exon 5, respectively (Fig. 1A), which result in complete loss of *DLDG1* mRNA synthesis (Supplementary Fig. S3). Both mutants had a pale-green phenotype under continuous light (Fig. 1D). Notably, the phenotype was observed only when the leaves were at the young stage, and the pale-green color disappeared by the time the leaves had matured (Fig. 1E). Interestingly, the pale-green phenotype was not observed when plants were grown under 12-h dark/12-h light conditions (Fig. 1F), suggesting that the negative impact on chloroplasts caused by *dldg1* mutation was restored during dark periods. To verify whether the delayed greening of *dldg1* leaves correlated with the length of the period kept in darkness, *dldg1-2* was grown under different light and dark periods. The percentage of plants showing delayed greening gradually decreased as the time kept in darkness increased, i.e., 100%, ~90%, ~75%, and 0% plants showed delayed greening when the dark period per 24 h was 0, 2, 4, and 8 h, respectively (Supplementary Table 1).

An analysis of cellular ultrastructure revealed that the chloroplasts in the pale-green leaves of *dldg1-1* were much smaller than those of WT (Fig. 1G). Furthermore, proper thylakoid membrane stacking was not observed in the pale-green leaves. The abnormal phenotypes were not observed in chloroplasts in the mature green leaves of *dldg1-1*.

The amounts of several chloroplast proteins were assessed by western blotting. The levels of the PSII subunits D1 and CP47 and the PSI subunit PsaA in the pale-green young leaves of *dldg1-1* that had been grown under continuous light were greatly decreased compared with those of WT (Fig. 2A). The decreases in the levels of CP47 and PsaA were fully and partially recovered, respectively, by the time the leaves had matured, although the D1 level was still very low compared with that of WT at the same age. The levels of the light-harvesting chlorophyll-binding protein of PSII, LHCB6, a subunit of the cytochrome *b*_*6*_*f* complex, PetC, and the ATP synthase subunit CF1γ, in *dldg1-1* were the same as those of WT in both young and mature leaves when grown under continuous light (Fig. 2A).

**Fig. 2.**
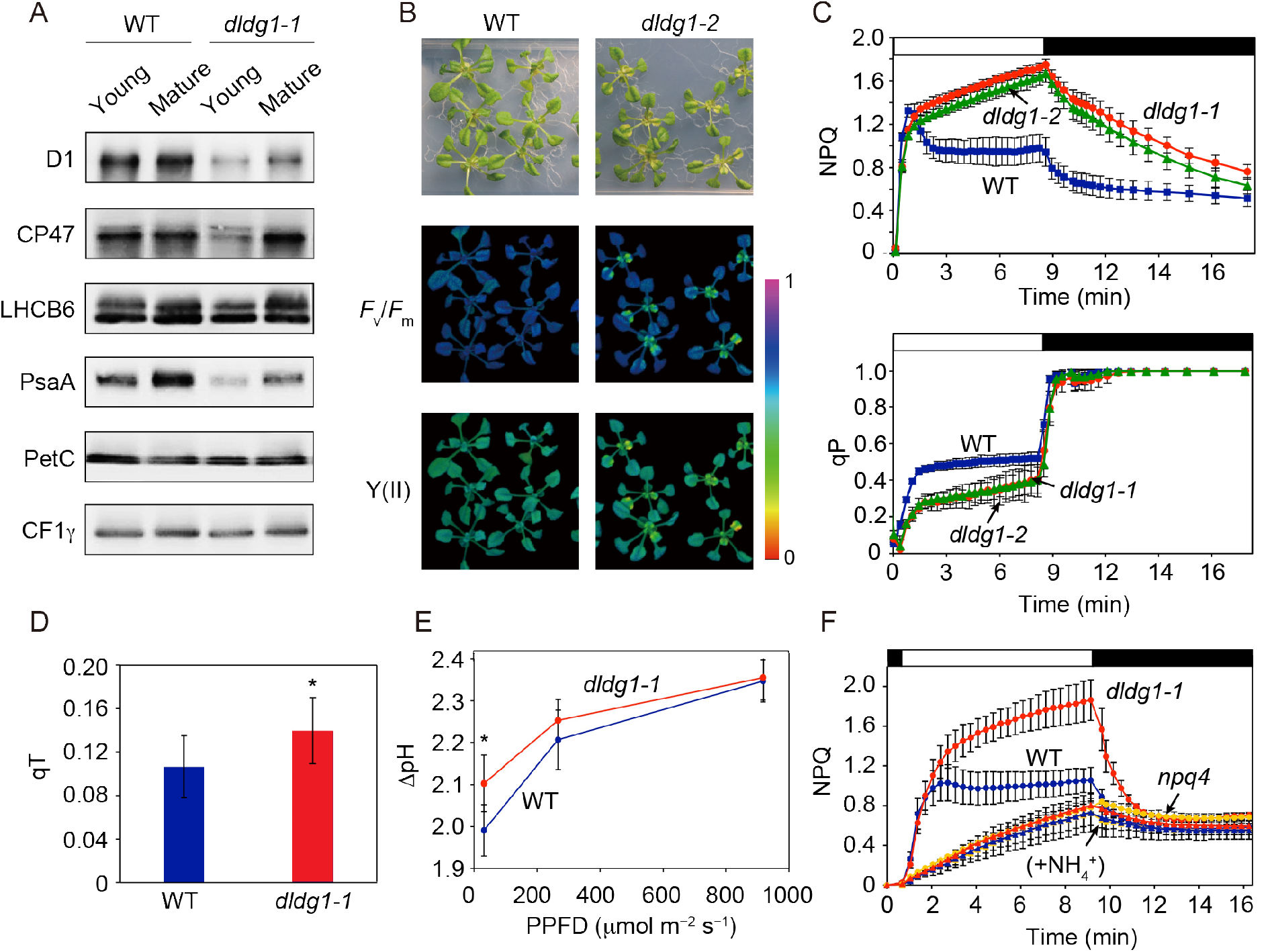
Protein levels and photosynthetic activities of *dldg1*. (A) Western blots of chloroplast membrane proteins described in the text. The same amounts of proteins were loaded onto each lane. (B) Images of WT and *dldg1-2* leaves used to assess the maximum PSII quantum yield (*F*_v_/*F*_m_) and the PSII quantum yield (Y(II)). Plants were grown on agar containing 1/2MS medium under continuous light (50 μmol m^−2^ s^−1^). Plants were dark-adapted for 30 min before measurements were made. For Y(II) measurements, the plants were illuminated with actinic light (55 μmol m^−2^ s^−1^) for 10 min. (C) Dark-to-light (230 μmol m^−2^ s^−1^) induced kinetics for NPQ and qP. The values are the mean ± SD (*n* = 5). (D) Quenching of chlorophyll fluorescence due to state transitions (qT). The values are the mean ± SD (*n* = 8; \**P* < 0.05). (E) Effect of light intensity on ΔpH in isolated chloroplasts of WT and *dldg1-1*. The values are the mean ± SD (*n* = 3; \**P* < 0.05). (F) Dark-to-light (230 μmol m^−2^ s^−1^) induced kinetics for NPQ of WT (blue), *dldg1-1* (red) and *npq4* (yellow) that were pre-incubated with 100 mM KCl (circles) or NH_4_Cl (triangles) for 30 min. The values are the mean ± SD (*n* = 5).

### Photosynthetic parameters of *dldg1*

The photosynthetic parameters of *dldg1* were measured using a pulse-amplitude modulation Chl fluorometer. Pale-green leaves from *dldg1-2* that had been grown under continuous light (50 μmol photons m^−2^ s^−1^) had a reduced maximum PSII quantum yield (*F*_v_/*F*_m_) and a reduced PSII quantum yield (Y(II)) compared with WT (Fig. 2B). In the mature leaves of *dldg1-2*, their *F*_v_/*F*_m_ and Y(II) levels were comparable to those of WT. We also measured the NPQ and qP induction kinetics, which represent the extent of non-photochemical quenching and oxidized Q_A_ in PSII, respectively. For this measurement, mature leaves that did not have a pale-green phenotype were used. WT and two *dldg1* plants showed transient NPQ induction after illumination by a photosynthetic photon-flux density (PPFD) of 230 μmol photons m^−2^ s^−1^ (Fig. 2C). By ~1 min after the start of illumination, the NPQ in the WT plants had relaxed to a lesser extent, although the two *dldg1* mutants sustained NPQ induction. After removing the actinic-light, a slower NPQ relaxation was observed in the *dldg1* mutants compared with in WT. The qP values in the two *dldg1* mutants were lower than in WT upon illumination, indicating that plastoquinone pool in *dldg1* tends to be reduced compared to those in WT. The qP values for WT and the *dldg1* mutants were the same immediately after the actinic-light illumination was turned off. NPQ induction kinetics was also measured with plants that were pre-treated with an ionophore NH_4_^+^ that deforms ΔpH across the thylakoid membranes (Fig. 2F). Both WT and *dldg1-1* showed significant reduction of NPQ induction after NH_4_^+^-treatment, and the kinetics were similar to those of NH_4_^+^-treated and untreated *npq4* mutant, indicating that the increment of NPQ in *dldg1* results from enhancement of the PsbS-dependent qE. We also assessed effect of *dldg1* mutation on qT quenching. qT induction is slightly, but significantly, higher in *dldg1-1* than in WT (Fig. 2D). qT is induced through LHCII phosphorylation by a protein kinase STN7, whose activity is up-regulated by reduction of plastoquinone pool (Dietzel et al., 2008), further suggesting that plastoquinone pool tends to be more reduced in *dldg1* than in WT.

Given PxcA, a DLDG1 homolog of cyanobacteria, is crucial for light-induced H^+^ extrusion, DLDG1 may also control H^+^ homeostasis in chloroplasts. To test this hypothesis, we measured internal pH change of isolated chloroplasts using 9-aminoacridine (9-AA) fluorometry. At neutral pH, 9-AA is uncharged and, therefore, has high membrane permeability. Once it protonated in weakly acidic solution, its membrane permeability as well as fluorescence yield is markedly reduced. Using this property, a system for measuring the internal pH of chloroplasts was established (Schuldiner et al., 1972). We used this system to measure ΔpH across the chloroplast envelope membrane under different light intensity. We isolated intact chloroplasts from plants grown under an 8-h light/16-h dark cycle as described previously (Kawashima et al., 2017), for which *dldg1* did not show the pale-green phenotype (Supplementary Table 1). The ΔpH value is significantly higher in *dldg1-1* than in WT under low-light (31 μmol photons m^−2^ s^−1^) conditions (Fig. 2E), indicating that internal pH (determined by H^+^ concentrations in both the lumen and stroma) is lower in *dldg1* than in WT at the condition. On the other hand, no difference was observed under middle- and high-light intensity (>~200 μmol photons m^−2^ s^−1^).

We next recorded electrochromic absorbance shift (ECS) signals to calculate the proton motive force (*pmf*), proton concentration gradient (ΔpH_*pmf*_) and electric potential gradient (ΔΨ_*pmf*_) across the thylakoid membranes, as well as thylakoid proton conductivity by ATP synthesis (g_H_^+^). For these measurements, we grew plants under an 8-h light/16-h dark cycle, for which *dldg1* did not show the pale-green phenotype. No significant difference was observed between the *pmf*, ΔpH_*pmf*_, and ΔΨ_*pmf*_ values of WT and the *dldg1-1* mutant under all PPFDs tested (Fig. 3A-C). Conversely, the average g_H_^+^ value for the mutant was significantly smaller than that of WT under a PPFD of ~100 μmol photons m^−2^ s^−1^ (Fig. 3D).

**Fig. 3.**
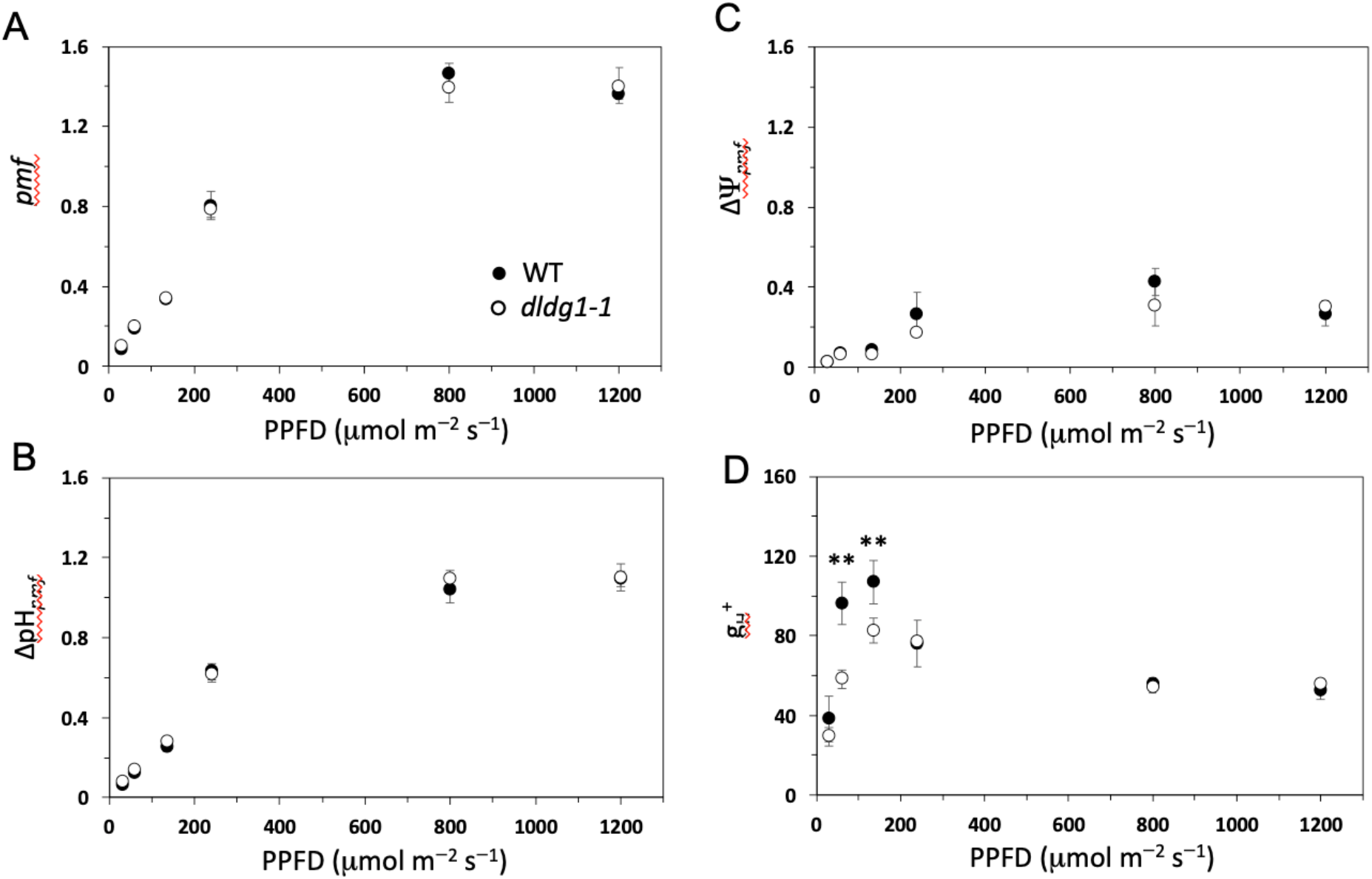
*pmf* responses to various PPFDs. (A) Total *pmf* vs. PPFD, (B) ∆pH_*pmf*_ vs. PPFD, (C) ∆Ψ_*pmf*_ vs. PPFD, and (D) proton conductivity (g_H_^+^) vs. PPFD, as estimated from the dark-interval relaxation kinetics of the ECS signal measured in the leaves of WT (●) and *dldg1-1* (○). The values represent the mean ± SD (*n* = 3–6; \*\**P* < 0.01).

Next, we measured Y(I), Y(ND), and Y(NA), which report values for the quantum yield, donor side-limited heat dissipation, and accepter side-limited heat dissipation of PSI, respectively (Fig. 4A). The measurements were made under illumination that fluctuated between a PPFD of 800 μmol photons m^−2^ s^−1^ for 10 min and 30 μmol photons m^−2^ s^−1^ for 15 min. The Y(ND) values for *dldg1-1* were greater than for WT starting with the second round of the two-step illumination (Fig. 4A, middle panel), although such a significant effect on the Y(I) and Y(NA) was not seen (top and bottom panels). We also measured NPQ and Y(II) under illumination that fluctuated between a PPFD of 91 μmol photons m^−2^ s^−1^ and 921 μmol photons m^−2^ s^−1^ (Fig. 4B). The NPQ values for *dldg1-1* were greater than for WT in both low and high PPFDs; the large NPQ phenotype of *dldg1-1* is more clearly observed under the high PPFD (Fig. 4B). Y(II) was also lowered in *dldg1-1* under the high PPFD, especially with the second round of the two-step illumination, indicating that negative impacts on photosynthesis regulation caused by the *dldg1* mutation appear strongly under fluctuating light.

**Fig. 4.**
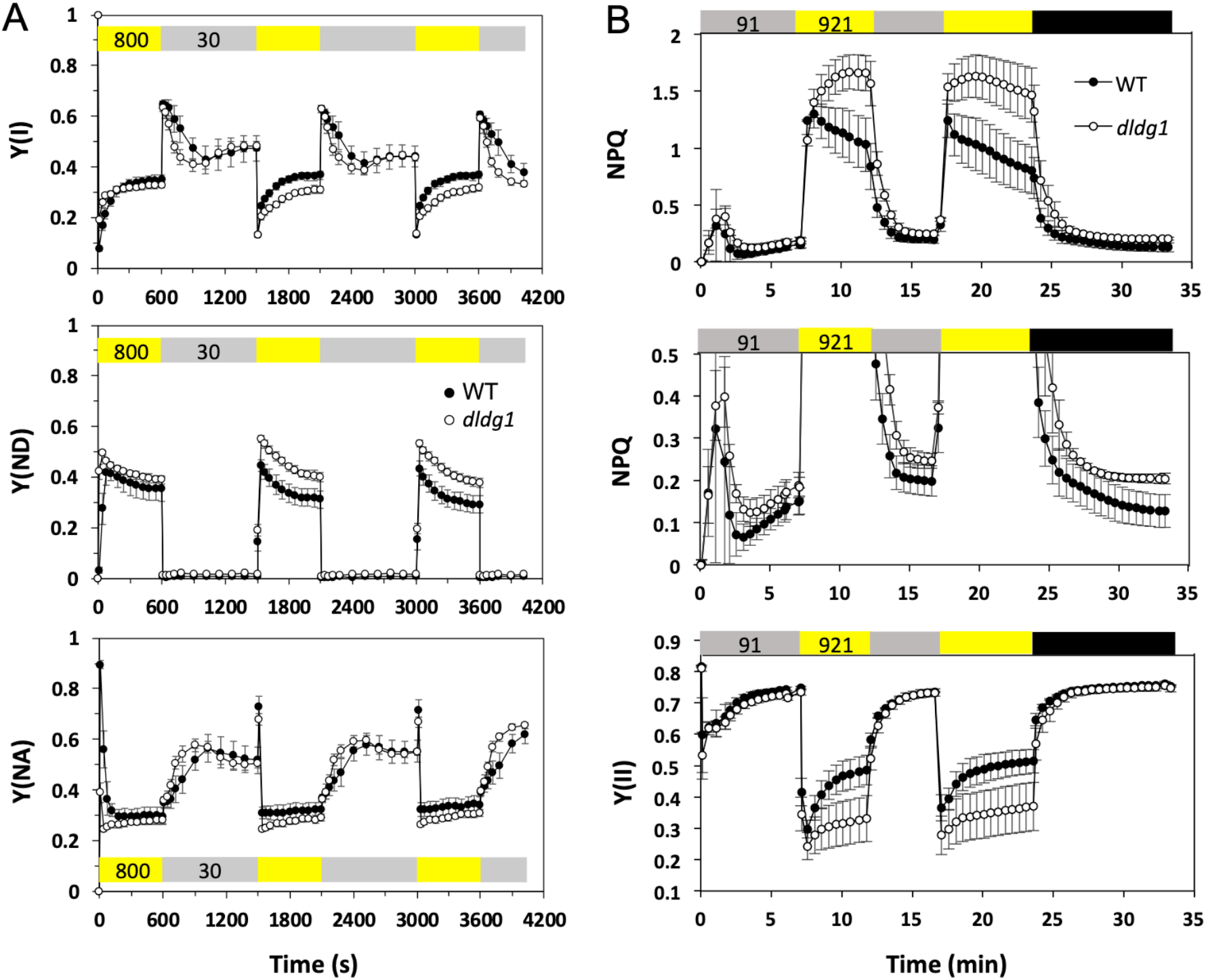
PSI and PSII activity under fluctuating light. PSI (A) and PSII (B) activity measurement. NPQ parameters shown in the top panel is expanded in the middle panel. The parameters were monitored under fluctuating light in the leaves of WT (●) and *dldg1-1* (○) plants. The light fluctuated between 800 μmol m^−2^ s^−1^ (for 600 sec) and 30 μmol m^−2^ s^−1^ (for 900 sec) for PSI activity measurement (a), and 91 μmol m^−2^ s^−1^ (for 7 min and 5 min at the first and the second round, respectively) and 921 μmol m^−2^ s^−1^ (for 5 min and 7 min at the first and the second round, respectively) for PSII activity measurement (b). Each data point represents the mean ± SD (*n* = 3–5).

We next characterized the carotenoid compositions of WT and *dldg1-1* (grown under the 8-h light/16-h dark cycle) before and after exposure to intense light (1,900 μmol photons m^−2^ s^−1^), which causes the accumulation of zeaxanthin for qE (or qZ) induction (Nilkens et al., 2010; Ruban, 2016). For *dldg1-1*, we found that the zeaxanthin content after exposure to intense light and the lutein content before and after exposure to intense light were significantly greater than in WT (~1.3- and ~1.2-fold, respectively; Supplementary Table 2). No significant difference was observed for the amount of any other carotenoid compound.

### Characterization of DLDG1 function

The *pxcA*-less mutant of the cyanobacterium *Synechocystis* sp. PCC6803 showed no light-induced H^+^ extrusion activity (Katoh et al., 1996), suggesting the function of PxcA-like proteins (such as DLDG1) to be related to ion transport across membranes. To test this hypothesis, we performed complementation analysis of the K^+^ uptake deficient *E. coli* strain LB2003 (Stumpe et al., 1996), with Arabidopsis DLDG1, Ycf10, and/or FLAP1. The LB2003 strain, containing the vector only, showed slow growth under low K^+^ concentrations, because it lacks four K^+^ uptake transporters (Fig. 5A). We found that complementing strains expressing DLDG1 showed more strong growth retardation than the vector-containing LB2003 (Fig. 5A), suggesting that DLDG1 has the K^+^ efflux activity. We also performed complementation analysis of the *E. coli* strain TO114, lacking three Na^+^/H^+^ antiporters (Ohyama et al., 1994). We used the Na^+^/H^+^ antiporter NhaS3 of *Synechocystis* as a positive control (Tsunekawa et al., 2009). As shown in Fig. 5B, the TO114 strain, containing vector(s) only, showed inability to grow in the presence of elevated concentrations of Na^+^, because of its lowered Na^+^ efflux/H^+^ influx activity (Ohyama et al., 1994), and the phenotype could be partially complemented when DLDG1 or Ycf10 was introduced, suggesting that DLDG1 and Ycf10 has the Na^+^ efflux activity.

**Fig. 5.**
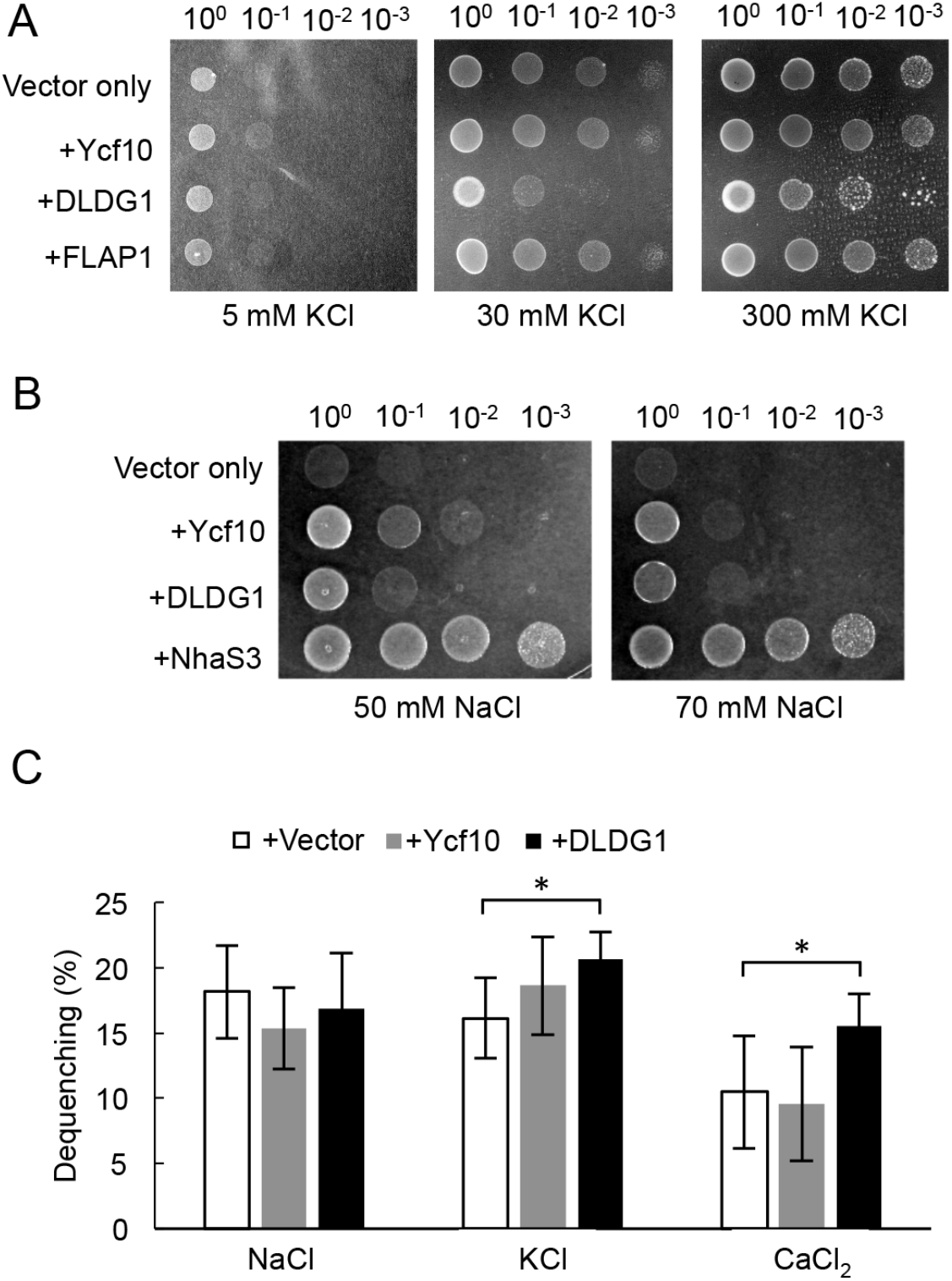
Transporter activity of DLDG1. (A) Complementation tests of *E. coli* LB2003 strain transformed with genes indicated. Cells were grown in the presence of 5, 30 and 300 mM KCl for 6, 4, and 3 days, respectively. (B) Complementation tests of *E. coli* TO114 strain transformed with genes indicated. Cells were grown in the presence of 50 and 70 mM NaCl for 1 day. (C) Antiporter activities in everted membrane vesicles of *E. coli* TO114 strain transformed with empty vectors, *DLDG1* and/or *Ycf10*. The values represent the mean ± SD (*n* = 3; \**P* < 0.05).

Antiporter activities were further analyzed by everted membrane vesicles of the transformed *E. coli*. The everted membrane vesicles provided from the *E. coli* expressing NhaS3 showed strong H^+^ extrusion activity upon Na^+^ addition; however, those of DLDG1 and/or Ycf10-expressing *E. coli* did not (Fig. 5C, Supplemental Fig. S4), suggesting that expressed DLDG1 and Ycf10 showed no or very low Na^+^/H^+^ antiport activity in the everted vesicles. However, membrane vesicles from the *E. coli* expressing DLDG1 showed small but significant higher K^+^/H^+^ and Ca^2+^/H^+^ antiport activities. Given the experiment was performed with everted vesicles, the result suggests that DLDG1 has a K^+^ (or Ca^2+^) efflux/ H^+^ import activity in native cells.

## Discussion

Our results indicate that Arabidopsis DLDG1, a plant homolog of the plasma membrane-localized cyanobacterial protein PxcA, is required for proper control of chloroplast development and NPQ induction. PxcA is necessary for light-induced H^+^ extrusion by cyanobacteria (Katoh et al., 1996; Sonoda et al., 1998). Specifically, illumination of dark-adapted WT *Synechocystis* cells results in an extrusion of H^+^ into the medium followed by H^+^ re-uptake; however, the *pxcA* mutant of *Synechocystis* lacks the ability to extrude H^+^, although proper H^+^-re-uptake activity is retained (Sonoda et al., 1998). Our computer-aided homology search indicated that there are two types of PxcA homologs in cyanobacteria and plants, i.e., nuclear-encoded DLDG1s and plastid-encoded Ycf10s in plants (Supplementary Fig. S2). Ycf10 orthologs, encoded in the plastid genomes of pea and the green alga *Chlamydomonas*, have been characterized (Sasaki et al., 1993; Rolland et al., 1997). Pea Ycf10 is localized in the chloroplast envelope membrane, and thus the protein was designated as <u>c</u>hloroplast <u>e</u>nvelope <u>m</u>embrane protein A (CemA) (Sasaki et al., 1993). Given that Arabidopsis DLDG1 also localizes in the envelope membrane of chloroplasts (Fig. 1C), these two proteins are functionally similar. The *Chlamydomonas* Ycf10-deficient mutant showed retarded growth under intense light, which may have been caused by a decreased uptake of inorganic carbon (Rolland et al., 1997). Ycf10 was suggested to be involved in pH regulation in chloroplasts by exporting reducing power into the cytoplasm and, in particular, by extrusion of H^+^ from the chloroplast stroma to the cytoplasm; such dynamic changes in H^+^ levels may cause a localized pH change near the intermembrane space of the chloroplast envelope and thereby stimulate conversion of HCO_3_^−^ into CO_2_ and vice versa, which would affect inorganic carbon uptake (Rolland et al., 1997). Here, we found that DLDG1 has the K^+^ efflux/H^+^ influx activity (Fig. 5), suggesting that it imports H^+^ from the cytoplasm to the chloroplast stroma. Based on these assumptions, we propose that Ycf10 and DLDG1 coordinately control the stromal pH by the H^+^ extrusion and import activities across the envelope membranes, respectively. This control may be required for efficient CO_2_ fixation, because Calvin-Benson cycle enzymes are sensitive to pH change (Werdan et al., 1975). Hence, inhibition of CO_2_ fixation accompanies an imbalance in the consumption of ATP and NADPH. The Arabidopsis *inap1* mutant has an unbalanced NADP status and exhibits a pale-green phenotype (Werdan et al., 1975) as does *dldg1* (Fig. 2D), which supports the hypothesis that DLDG1 and Ycf10 control H^+^ homeostasis in the chloroplast stroma and thereby modulates the reducing power provided by photosynthesis. In fact, NPQ mutants such as *npq4* have no growth or developmental phenotype (Li et al., 2000), indicating that the impaired regulation of NPQ cannot be behind the mutant *dldg1* phenotype.

Strict coordination of stroma pH may also involve lumen pH regulation because water oxidation in the PSII continuously produces H^+^ that may be transferred from the lumen into the stroma by the ATP synthase, membrane transporters and/or ion channels whose activity might be influenced by the stromal pH change. Perhaps, in the *dldg1* mutant, lumen pH is lowered than in WT, since acidification of the lumen is known to be accompanied enhanced protonation of PsbS and VDE that results in a larger NPQ (Fig. 2C, Fig. 4B) and zeaxanthin accumulation (Supplementary Table 2). Acidification of the lumen is accompanied by enhanced protonation of PsbS and VDE that results in a larger NPQ (Fig. 2C, Fig. 4B) and zeaxanthin accumulation (Supplementary Table 2). Lumen acidification also affects feedback control of other critical steps caused by light-induced reactions including: 1) prevention of *b*_*6*_*f* complex activity to suppress electron accumulation on PSI (Werdan et al., 1975), 2) release of Ca^2+^ from the oxygen-evolving complex of PSII to limit electron flow by slowing the oxygen-evolving complex–dependent water oxidation (Semin et al., 1998), and 3) reduction of lumen-localized enzyme activities (Semin et al., 1998). All or at least some of these reactions may have been affected by lumen acidification in *dldg1* leaves that was the result of an imbalance in the redox state of the quinone pool with an unusual qP (Fig. 2C), qT (Fig. 2D), and Y(ND) (Fig. 4A) as well as a reduction in D1 levels with small *F*_v_/*F*_m_ values especially (Fig. 2A, B), which might contribute to its pale-green phenotype and unusual chloroplast ultrastructure (Fig. 1D, G). Given that these *dldg1* phenotypes were moderated when the plants were grown under a light/dark cycle (Fig. 1F), it appears that induction of stress in *dldg1* requires incident light, which further supports the hypothesis that DLDG1 is involved in the control of H^+^ homeostasis in chloroplasts.

Although the internal pH in chloroplasts were lower in *dldg1* than in WT, especially under low PPFD (Fig. 2E), *pmf*, ΔpH_*pmf*_, and ΔΨ_*pmf*_ values were similar in WT and *dldg1* (Fig. 3A-C), suggesting that acidification, caused by the *dldg1* mutation, is simultaneously altered to the same extent in both the lumen and stroma. In WT, g_H_^+^ increased by ~two-fold under moderate light intensity (~100 μmol photons m^−2^ s^−1^) compared with the values under low PPFD (<50 μmol photons m^−2^ s^−1^) or intense PPFD (>150 μmol photons m^−2^ s^−1^), as reported(Avenson et al., 2005); however, the increases in g_H_^+^ in *dldg1* were less than in WT (only ~1.2-fold increase; Fig. 3D). The aforementioned results suggest that the enhancement of ATP synthase activity induced by moderate light is countered by an increase in the acidity in the lumen and/or stroma of *dldg1*. Notably, the Arabidopsis mutant that lacks the chloroplast ATP synthase γ subunit exhibits destabilized ATP synthase activity and has decreased *DLDG1* expression (Bosco et al., 2004), suggesting that regulation of DLDG1 activity is coupled to ATP synthase activity.

Although expressed DLDG1 and/or Ycf10 complement or enhance the phenotype of the *E. coli* Na^+^/H^+^ and K^+^/H^+^ antiporter mutants (Fig. 5A, B), exact activities of the two proteins were still not fully clarified. The everted membrane vesicles of the *E. coli* expressing DLDG1 showed low, but significant activities of not only K^+^/H^+^ antiport but also Ca^2+^/H^+^ antiport, suggesting that the substrate selectivity of the counter ion(s) during H^+^ transport is low. Furthermore, everted membrane vesicles of the *E. coli* expressing Ycf10 did show antiport activities (Fig. 5C), although it can complement the phenotype of the *E. coli* Na^+^/H^+^ mutant (Fig. 5B). One possibility is that H^+^ efflux and influx by the Ycf10 orthologs require other components not identified yet, although no direct evidence has been provided.

We observed a slight, but significant, greater accumulation of lutein in *dldg1* than in WT (Supplementary Table 2). Lutein accumulation has been shown to complement NPQ induction in the zeaxanthin-less Arabidopsis *npq1* mutant (Li et al., 2009; Dall’Osto et al., 2017), suggesting that the accumulated lutein contributes to the increased rate of NPQ of *dldg1*, which may be caused by feedback control through acidification of the lumen and/or stroma, although the mechanism remains to be delineated.

A comprehensive yeast two-hybrid analysis was previously carried out with all open-reading-frames found in *Synechocystis* sp. PCC6803 genome; the results indicated that PxcA interacts with the cyanobacterial FLAP1 homolog, slr0404 (Sato et al., 2007). We reported that Arabidopsis FLAP1 is found in the thylakoid and envelope membrane of chloroplasts and that a *flap1* mutant had a greater NPQ value than WT, especially under fluctuating light (Sato et al., 2017; Trinh et al., 2019); the phenotype is similar to those observed in *dldg1* (Fig. 4B). These results suggest that FLAP1, DLDG1, and Ycf10 orthologs, which localize in chloroplast envelope or cyanobacterial plasma membranes, physiologically interact to function, although their exact activities have not been fully delineated. Conversely, thylakoid membrane–localized FLAP1 may potentially interact functionally with as-yet unidentified partners other than DLDG1/Ycf10. Recently, several ion transporters, localized in thylakoid membranes, were identified that regulate photosynthetic output and NPQ (Szabò and Spetea, 2017). One possibility is that FLAP1 functionally interacts with such transporters. Clearly, further characterization of multiple mutants of FLAP1, DLDG1, and Ycf10 orthologs in cyanobacteria and plants will be needed to further characterize the mechanisms and evolution of H^+^/ion homeostasis control and its physiological importance for regulating oxygenic photosynthesis.

## Materials and Methods

### Plants and growth conditions

Plants were the Columbia-0 ecotype of *A. thaliana* and were grown at 23°C on 0.8% (w/v) agar containing half-strength Murashige and Skoog (1/2MS) medium or in soil containing 50% (w/w) vermiculite. Plants were grown under different light/dark-period and light-intensity conditions, which are indicated in each figure legend and/or experimental section. The T-DNA insertional Arabidopsis *dldg1-1* (WiscDsLox413-416O18) and *dldg1-2* (SAIL-253-F06) mutants were obtained from the Arabidopsis Biological Resource Center (Columbus, OH). Using gene-specific primers, homozygous insertion of each *dldg1* mutation was confirmed by PCR followed by sequencing.

### Reverse-transcription PCR

Total RNA was isolated using the SV Total RNA Isolation System (Promega). First-strand cDNA, synthesized using the PrimeScript RT regent Kit (TaKARa), or isolated gDNA was the subsequent template. Primers A, B, C, and D (Supplementary Table S3) were used specific primers for *DLDG1* amplification, and those for *CRSH* were as described previously(Maekawa et al., 2015).

### Measuring Chl fluorescence

Plants were grown in 1/2 MS medium under continuous light, and the induction kinetics of Chl fluorescence in rosette leaves, which did not have the pale-green phenotype, of 21-day-old plants were recorded using an IMAGING-PAM Maxi Version Chlorophyll Fluorometer (Walz, Effeltrich, Germany). Before each measurement, plants were adapted to the dark for 15 min. The minimum Chl fluorescence at an open PSII center (*F*_*o*_) was measured with excitation by light at 655 nm (0.05–0.15 μmol photons m^−2^ s^−1^). A saturating light pulse was applied to determine the maximum Chl fluorescence at the closed PSII centers when in the dark (*F*_*m*_) and during illumination with actinic light (*F*_*m*_*′*). The steady-state Chl fluorescence (*F*_*s*_) was recorded during continuous illumination with actinic light. The maximum quantum efficiency (*F*_*v*_/*F*_*m*_) was calculated as (*F*_*m*_ – *F*_*o*_)/*F*_*m*_. The operating efficiency, Y(II), during steady-state photosynthesis was calculated as (*F*_*m*_′ – *F*_*s*_)/*F*_*m*_′. NPQ was calculated as (*F*_*m*_ – *F*_*m*_′)/*F*_*m*_′. qP was calculated as (*F*_*m*_′ – *F*_*s*_)/*F*_*m*_′ – *F*_*o*_′).

NPQ measurement with fluctuating actinic-light as well as qT analysis were carried out using Dual PAM-100 (Waltz) with plants grown under a 16-hr dark/8-hr light (200 μmol m^−2^ s^−1^) cycle for 50-60 days. qT calculations were performed as described(Lunde et al., 2000). Preferential PSII excitation was provided by red light (31 μmol m^−2^ s^−1^) (State 2 light). For transition to state 1, leaves were exposed to the red light supplemented with far-red light (State 1 light). qT parameters were calculated as (*Fm*_*1*_ − *Fm*_*2*_)/*Fm*_*1*_, where *Fm*_*1*_ and *Fm*_*2*_ designate the maximal fluorescence yield in State 1 and State 2, respectively. 9-AA quenching analysis was performed as described(Kawashima et al., 2017).

### Measurement of the ECS signal

Plants were pot-grown in a growth chamber at 23°C and relative humidity 60%. During the 8-h photoperiod, light was provided by a bank of white fluorescent tubes, and the irradiance at the plant level was 135 μmol m^−2^ s^−1^. Plants were irrigated two to three times a week with deionized water for 2 weeks after germination, and afterwards with a 1:500 strength Hyponex 6-10-5 solution (Hyponex Japan). Mature rosette leaves from 8- to 9-week-old plants were used in experiments.

The *pmf* was assessed by measuring the ECS with a DUAL-PAM-100 instrument containing a P515/535 module. The ECS signal was the difference between the transmittance at 550 nm and that at 515 nm(Klughammer et al., 2013). DIRK-analysis (Baker et al., 2007) was conducted to determine the value of g_H_^+^ along with values for the proton conductivity of the thylakoid membrane(Joliot and Joliot, 2002; Avenson et al., 2005) and the two *pmf* components, ∆pH_*pmf*_ and ∆Ψ_*pmf*_ (Sacksteder et al., 2000; Jeffrey A. Cruz et al., 2001). Leaves were each illuminated with red and blue light at 150 μmol m^−2^ s^−1^ for 30 min, and then the PPFD value was gradually increased to 1,200 μmol m^−2^ s^−1^ to open the stomata. Next, the blue light was removed and measurements were made. At each PPFD intensity, the DIRK of the ECS signal was obtained 300 s after the onset of the actinic light. Measurements were made in an air-ventilated room (40 Pa CO_2_, 21 kPa O_2_, 25°C). ECS signals were normalized to that of a single-turnover flash. The ECS decay was measured for 200 to 300 ms after actinic light was removed and was fit to the one-component decay kinetics equation, A_1_e^−k1t^ + B, where A_1_ is the amplitude constant, k_1_ is the rate constant, and B is a constant. g_H_^+^ were calculated as the inverse of a decay time constant for a 200 – 300 ms dark period, respectively.

### Changes in the absorbance at 830 nm

Plants were grown under the same conditions as for the ECS measurements described above. Absorption changes at 830 nm were measured simultaneously for an intact leaf in a Dual-PAM Gas-Exchange cuvette (3010-Dual; Walz) with a GFS-3000 Portable Gas Exchange System (Walz) and a DUAL-PAM-100 Chl fluorescence and P700 absorption analyzer equipped with a P700 dual-wavelength emitter (at 830 and 875 nm). The CO_2_ concentration in the leaf chamber was regulated by a Control Unit 3000-C in the GFS-3000 system. The concentration of O_2_ was held constant by mixing N_2(g)_ and O_2(g)_ using mass flow controllers. The CO_2_ and O_2_ partial pressures in the leaf chamber were 40 Pa and 20 kPa, respectively. The leaf temperature was 23°C. The vapor pressure deficit ranged from 0.6 to 0.75 kPa. P700^+^ was monitored as the absorption difference between the 830 and 875 nm transmission measurements. Before starting the measurements, plants had been held under a light of intensity135 μmol m^−2^ s^−1^ for 30 to 40 min to activate enzymes of the Calvin-Benson cycle and open the stomata. The plants were subsequently kept in the dark for 5 min. The leaf lamina was sandwiched in the chamber. Saturation pulses of far-red light at 740 nm from LEDs (7,000 μmol m^−2^ s^−1^; 400-ms duration) were applied to determine the maximum oxidizable P700 (P_m_ value). The PSI quantum yields were determined using the saturation-pulse method(Klughammer and Schreiber, 1994). Y(I), Y(ND), and Y(NA), which sum to one, were determined under actinic-light conditions. To oxidize the inter-system electron carriers, we applied far-red light from 200 ms before the start of the saturation pulse application to its cessation.

### Transmission electron microscopy

Areas of pale-green and non-pale-green rosette leaves were cut from 30-day-old plants grown on 1/2MS medium under continuous light. These leaf segments were fixed and embedded in Spurr’s low-viscosity resin as described (Maekawa et al., 2015). Thin sections were stained with uranyl acetate and lead citrate and visualized with a Hitachi H7500 transmission electron microscope operating at 80 kV.

### Transient expression of GFP-fusion proteins

The DNA fragment for the N-terminal 150 residues of DLDG1 (corresponding to the predicted plastidial transit peptide) was PCR-amplified with the use of primers DLDG1-N150-F and DLDG1-N150-R (Supplementary Table S3), and the amplified fragment was used to make an in-frame DLDG1_N150_-GFP fusion construct. The final plasmid construct, 35S:DLDG1_N150_-GFP-pUC18, and the empty vector, 35S:GFP-pUC18 (kindly provided by Dr. Niwa of University of Shizuoka), were transiently expressed in onion epidermal cells after being introduced with a particle gun (PDS-1000He Particle Delivery System, Bio-Rad). The bombarded cells were kept in the dark at 22°C for 16 h followed by GFP imaging using a fluorescent microscope (ECLIPSE 80i, Nikon).

Full-length *DLDG1* cDNA was amplified by reverse-transcription PCR with DLDG1-F and DLDG1-R primers (Supplementary Table S3), and the PCR product was cloned into pDONRzeo using BP Clonase II reagents (Invitrogen). The inserted fragment was then isolated and cloned into pGWB5 (kindly provided by Dr. Nakagawa of Shimane University) using LR Clonase II reagents (Invitrogen). The resultant construct, 35S:DLDG1-GFP-pGWB5, the empty vector, 35S:GFP-pGWB5, and 35S:SFR2-EYFP-pGWB441(Sato et al., 2017) were separately introduced into *Agrobacterium tumefaciens* cells by electroporation. The cells were cultured in Luria-Bertani medium until late-log phase and then pelleted by centrifugation at room temperature. The harvested cells were suspended in ~15 ml of 10 mM MES-KOH (pH 5.5), 10 mM MgSO_4_, and 2% (w/v) sucrose, and then the OD_600_ of the suspension was adjusted to ~0.7. Each suspension was syringe-injected into *Nicotiana benthamiana* leaves. Plants were incubated at 23°C for 3 days under continuous light (40 μmol photons m^−2^ s^−1^). Fluorescence from GFP, YFP, and Chl was observed with a confocal laser-scanning microscope (LSM780, ZEISS).

### Western blotting

Samples of thylakoid membrane proteins from WT and *dldg1-1* plants were subjected to SDS-PAGE, and the separated protein bands were electroblotted onto a polyvinylidene difluoride membrane (GE Healthcare). Different amount of proteins were used for each western blotting; 2 μg (for anti-D1), 3 μg (for anti-PsaA), 20 μg (for anti-CF1γ) or 30 μg (for anti-CP47, anti-LHCB6 and anti-PetC) of proteins were loaded in each lane. Immunodetection was carried out using specific antibodies and an ECL Plus Western Blotting Detection System (GE Healthcare).

### Measurements of antiport activity of Ycf10/DLDG1

Inverted membrane of *E. coli* transformants were prepared as described previously (Kuroda et al., 1994). The inverted membranes equal to 1.2 mg of protein were added to the assay mixture (10 mM Tris-HCl (pH7.0 or pH8.0), 140 mM choline chloride, 5 mM MgSO_4_ and 1 µM acridine orange. To initiate the fluorescence quenching, Tris-_DL_-Lactate was added to a to final concentration of 10 mM. Subsequently, NaCl was added to a final concentration of 5 mM to observe Na^+^/H^+^, K^+^/H^+^ and Ca^2+^/H^+^ antiport activity. 50 μM of carbonyl cyanide 3-chlorophenylhydrazone (final concentration) was added to collapse ∆pH across the inverted membrane. The fluorescence was monitored at 525 nm with excitation light at 492 nm. Typical traces of the measurement were shown in Supplementary Fig. S4.

### Complementation analysis of the antiporter mutants of *E. coli*

Cording regions for *DLDG1*, *Ycf10* and *FLAP1* of Arabidopsis were amplified by reverse-transcription PCR with primer pairs; pPAB-DLDG1-F/pPAB-DLDG1-R, pPAB-Ycf10-F/pPAB-Ycf10-R, and pPAB-FLAP1-F/pPAB-FLAP1-R, respectively (Supplementary Table S3), and the PCR products were separately cloned into the *Bam*HI-*Sal*I-cut pPAB404(Buurman et al., 1995), and then separately introduced into *E. coli* strain TO114 (Ohyama et al., 1994) and/or LB2003 (Stumpe et al., 1996). Similarly, cording regions for *Ycf10* and *FLAP1* of Arabidopsis were also amplified by reverse-transcription PCR with primer pairs; pSTV-Ycf10-F/pSTV-Ycf10-R, and pSTV-FLAP1-F/pSTV-FLAP1-R, respectively (Supplementary Table S3), and the PCR products were separately cloned into the *Eco*RI-*Pst*I-cut pSTV28-tetr-KtrA (Matsuda et al., 2004) (replace *ktrA* with the PCR product). Obtained plasmids were separately introduced into the TO114 or LB2003 strains containing the pPAB404:DLDG1 plasmid as described below. The pSTV28-tetr:Ycf10 was also introduced into the pPAB404:FLAP1 plasmid. The procedure for the complementation analysis was described previously (Matsuda et al., 2004).

### Phylogenetic analysis

The amino-acid sequences of DLDG1 homologs were aligned with MAFFT(Katoh and Standley, 2013). The phylogenetic tree was constructed with RAxML(Stamatakis, 2014) with 1,000 bootstrap steps.

### Extraction and analysis of carotenoids

Seeds from Arabidopsis WT and *dldg1-2* plants were sown on 0.8% (w/v) agar containing 1/2MS medium and incubated under constant light (23°C, 40 μmol photons m^−2^ s^−1^) for 10 days to produce seedlings that were then transferred into soil and grown for 36 days under a 16-h dark (16°C)/8-h light (23°C) cycle (40 μmol photons m^−2^ s^−1^). Before and after being exposed to intense light (1,900 μmol photons m^−2^ s^−1^ for 40 min), five leaves from each plant were harvested, put immediately into liquid nitrogen, and then stored at −80°C until used for carotenoid analysis, which followed a published procedure (Kuwabara et al., 1998) with slight modifications. Specifically, each harvested leaf sample, in a 2-ml microtube, was treated with a mixture of acetone and methanol (7:2, v/v) containing 10 mM Tris/HCl (pH 8.0) to extract all pigments by mildly shaking each tube for several minutes under dim light. Each pigment-containing solution was placed into a small glass tube and subjected to rotary evaporation at 40°C. A small volume of a solution containing chloroform and methanol (3:1, v/v) was then added into each tube to dissolve the pigments. The samples were held in the dark until HPLC was performed to avoid light-induced isomerization. Each sample (10–20 μl) was chromatographed through a NOVA pack C18 column (RCM-type, 8 × 100 mm, Waters) and eluted with a solution containing 94.75% (v/v) acetonitrile, 1.75% (v/v) methanol, 1.75% (v/v) dichloromethane, and 1.75% (v/v) water for 12 min and then a solution containing 50% (v/v) acetonitrile, and 50% (v/) ethyl acetate for 20 min. A photodiode array detector (Waters 2998) was used to monitor the absorbance of the effluent (between 250 and 700 nm at 1.2-nm intervals) to detect the eluted pigments. The relative amount of each carotenoid is reported as its peak area normalized to that of the Chl *a* peak area in the same sample. The experiment was repeated four independent times, and values are expressed as the mean ± SD.

## Abbreviations

CemA: Chloroplast envelop membrane protein A
Chl: chlorophyll
DLDG1: Day-length-dependent delayed-greening1
CYO1: Cotyledon-specific chloroplast biogenesis factor1
ΔpH: H^+^ gradient across the thylakoid membrane
ΔΨ: electric potential
EYFP: enhanced yellow fluorescent protein
FLAP1: Fluctuating-light acclimation protein1
GFP: green fluorescent protein
LHC: light-harvesting complexes
NPQ: non-photochemical quenching
PS: photosystem
*pmf*: proton motive force across the thylakoid membrane
PPFD: photosynthetic photon-flux density
PxcA: proton-extrusion-protein A
SFR2: Sensitive to freezing2
VDE: violaxanthin de-epoxidase

## Acknowledgments

We thank Dr. Maki Kondo (National Institute for Basic Biology, Japan) for performing the electron microscopy study, and Professor Toshiharu Shikanai, Professor Krishna K. Niyogi and Arabidopsis Biological Resource Center for providing mutant seeds.

---

**Fig. S1.**
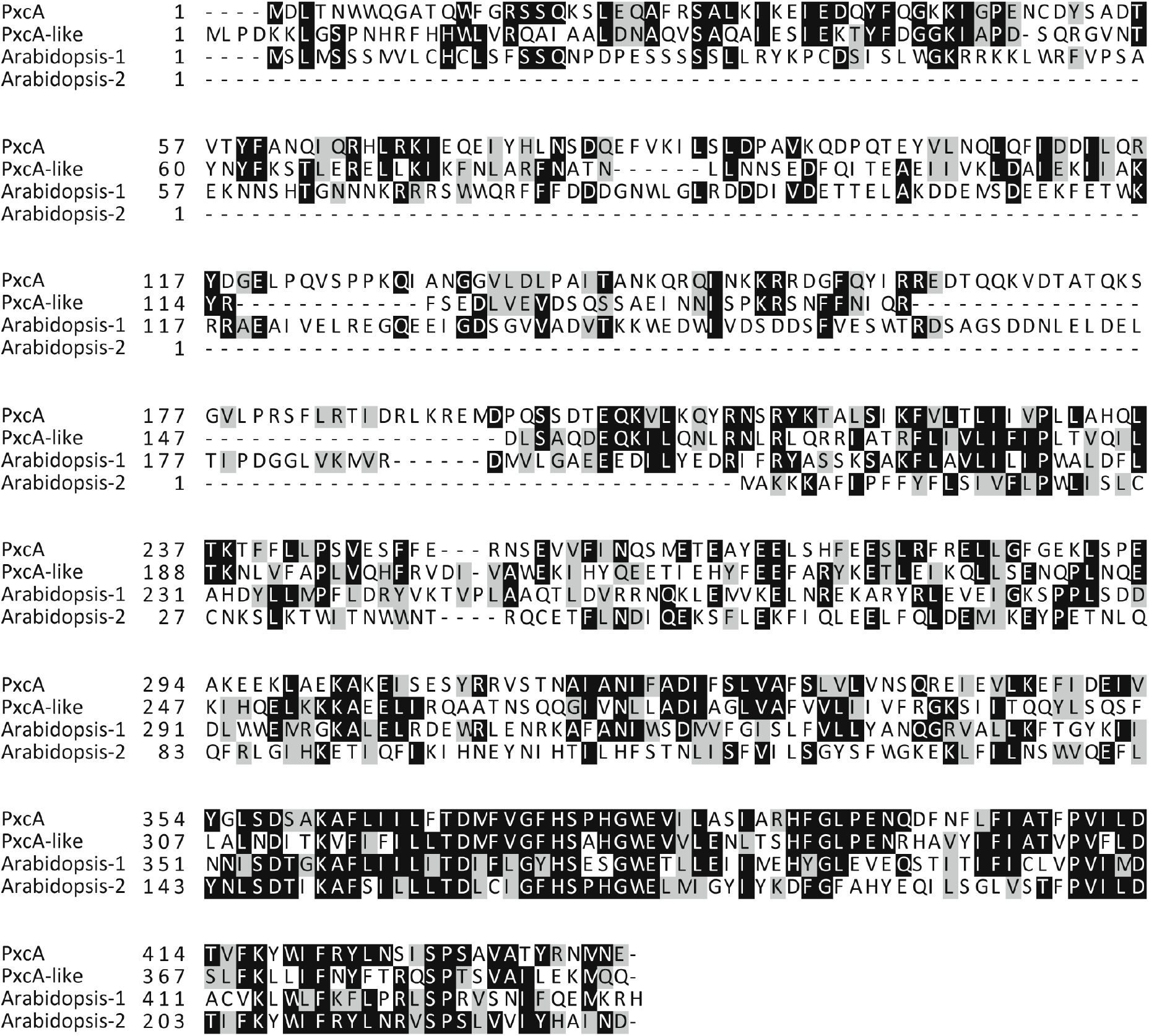
The amino-acid sequence alignment of DLDG1 homologs. Amino-acid sequences of PxcA (slr1596) and PxcA-like (sll1685) of the cyanobacterium *Synechocystis* sp. PCC6803 as well as Arabidopsis DLDG1 (Arabidopsis-1) and Ycf10 (Arabidopsis-2) are aligned.

**Fig. S2.**
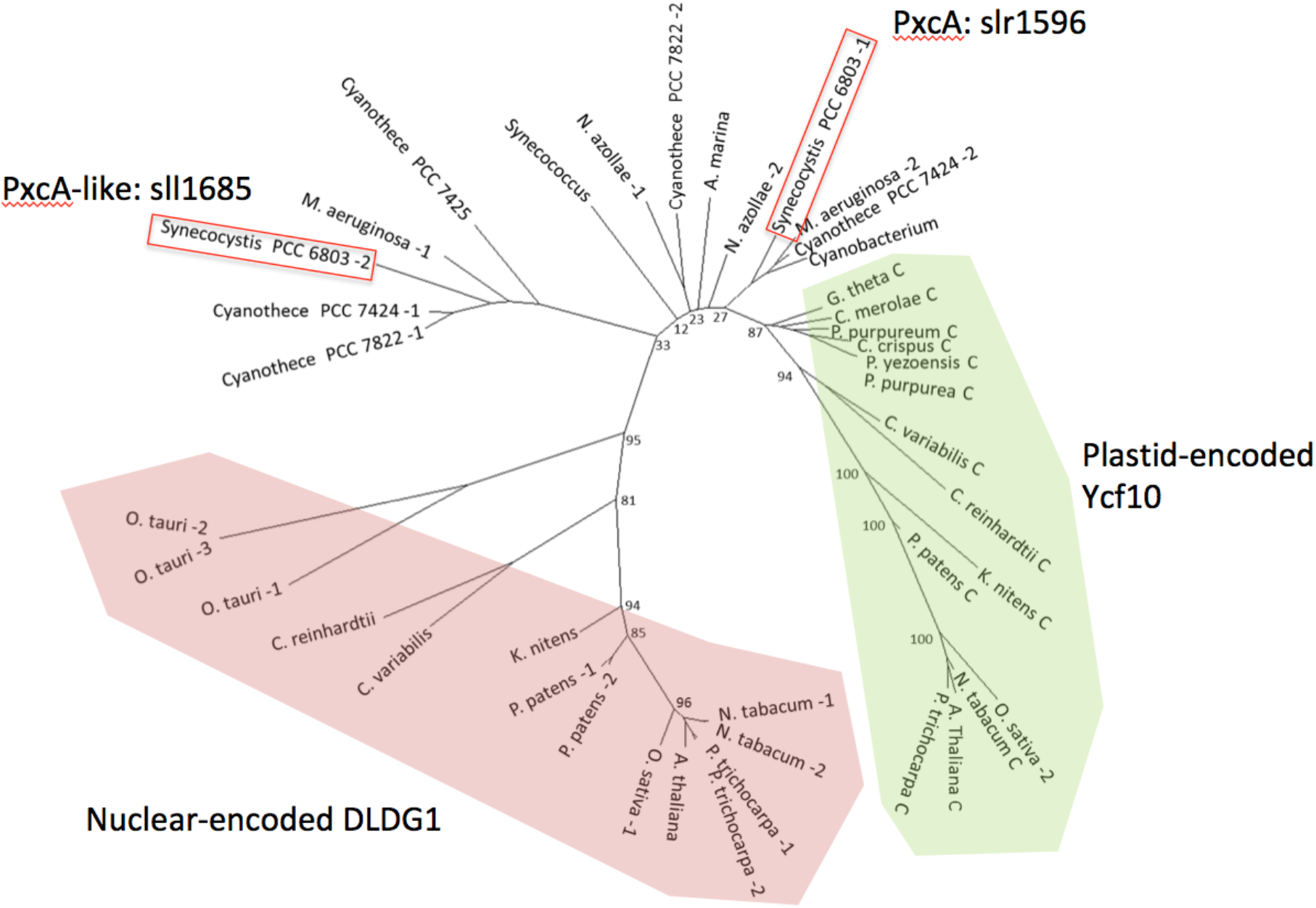
The phylogenetic tree of DLDG1 homologs. Plastid-encoded Ycf10 homologs and Nuclear-encoded DLDG1 homologs were highlighted by green and red, respectively. Two PxcA homologs of the cyanobacterium *Synechocystis* sp. PCC6803 are indicated by red squares. The sequence of *Populus trichocarpa* (1: XP_006372256, 2: XP_002309032.1, C: YP_001109514.1), *Oryza sativa* (1: XP_015648443.1, 2: NP_039398), *Nicotiana_tabacum* (1: XP_016433869.1, 2: XP_016440822.1, C: NP_054511.1), *Physcomitrella patens* (1: XP_024386612.1, 2: XP_024374481.1, C: NP_904190), *Klebsormidium nitens* (GAQ79310.1, C: GAQ93765.1), *Chlorella variabilis* (XP_005844579.1, C: YP_004347791.1), *Chlamydomonas reinhardtii* (XP_001696592.1, C: NP_958408.1), *Ostreococcus tauri* (1: XP_003078432.1, 2: XP_003080055.1, 3: XP_003080055.2), *Porphyra purpurea* (C: NP_053842.1), *Guillardia theta* (C: NP_050727.1), *Pyropia yezoensis* (C: YP_536913.1), *Chondrus crispus* (C: YP_007627444.1), *Porphyridium purpureum* (C: YP_008965641.1), *Cyanidioschyzon merolae* (C: NP_849136.1), *Cyanothece sp. PCC7822* (1: WP_013320969.1, 2: WP_013320969), *Cyanothece sp. PCC 7424* (1: WP_015954758.1, 2: WP_015955645.1), *Cyanothece sp. PCC 7425* (WP_012626986), *Microcystis aeruginosa* (1: WP_002762444, 2: WP_002787816.1), *Nostoc azollae* (1: WP_013189992.1, 2: WP_013191834.1), *Cyanobacterium* (WP_012954141.1), *Acaryochloris marina* (WP_012164771.1), *Synechococcus* (WP_011128082.1), were obtained from NCBI database using blastp program with sequence of *Arabidopsis thaliana* (NP_567865.1, C: NP_051071.1) and *Synechocystis PCC6803* (1: WP_010871416.1, 2: WP_010871621.1). The sequences were aligned using MAFTT 7.222 and trimmed by trimAl v1.2 with automated1 option to remove poorly aligned regions. RAxML (8.2.9, PROTGAMMALG, 1,000 bootstrap) were used for construction of phylogenetic tree.

**Fig. S3.**
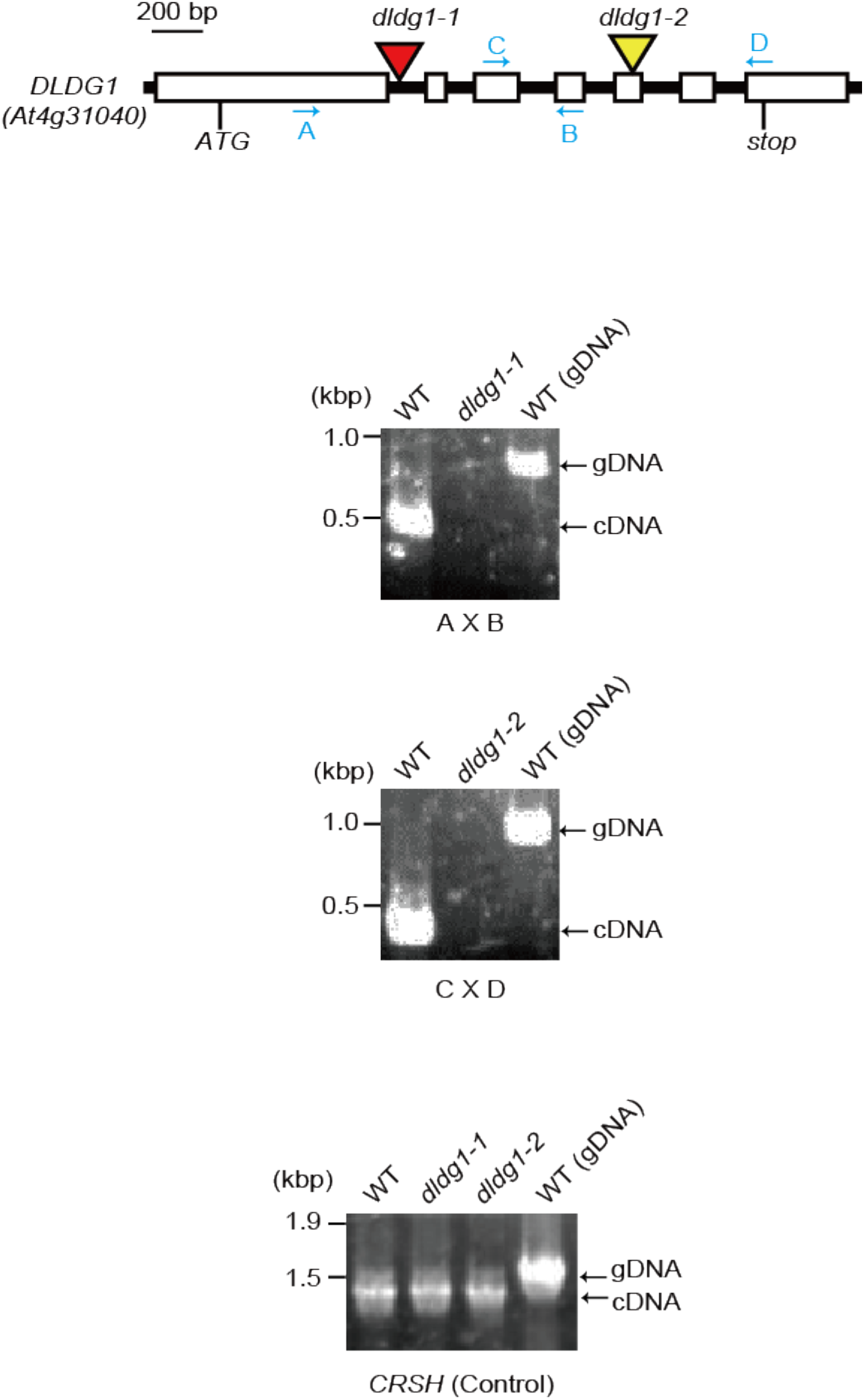
Two *dldg1* mutants showed no transcription of *DLDG1*. Positions of primers and T-DNA insertion sites are indicated. Reverse-transcription PCR were achieved by the primer pairs indicated at the bottom of each photo. *CRSH* was amplified as control for checking quality and quantity of RNA isolated. No amplification of genomic DNA (gDNA) was found for each PCR, indicating no contamination of gDNA in the RNA samples.

**Fig. S4.**
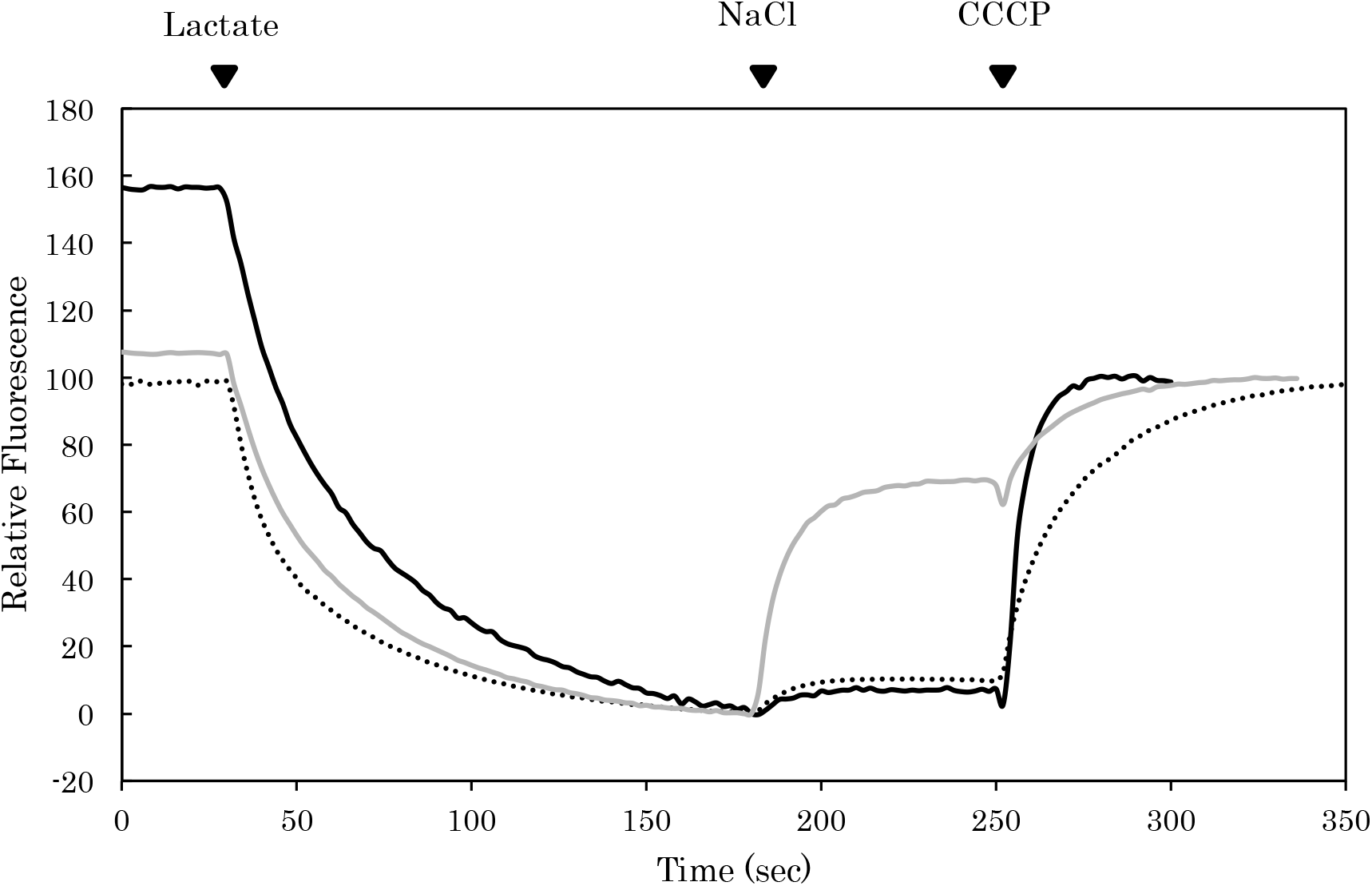
Typical traces for measuring the Na^+^/H^+^ antiport activity. Na^+^/H^+^ antiport activities of inverted membranes prepared from TO114 cells containing pPAB404/pSTV28-Tet^r^ (dotted line), pPAB404-DLDG1/pSTV28-Tet^r^-Ycf10 (black line) and pPAB404-NhaS3/pSTV28-Tet^r^ (gray line) were analyzed. 10 mM of Tris-DL-lactate was added to initiate quenching of fluorescence by respiration. 5 μM of NaCl was added to analyze Na^+^/H^+^ antiport activity. 50 μM of carbonyl cyanide 3-chlorophenylhydrazone (CCCP) was added to collapse ∆pH across inverted membranes.

**Table S1.** Percentage of individuals showing delayed greening phenotype under different light-dark periods. 40-50 individuals of WT and *dldg1-1* were grown on 1/2MS plates under different dark-light cycles as indicated. After 22 days, individuals showing pale-green phenotype were counted and the results are expressed as percentage.

|  | 24 h light/<br>0 h dark | 22 h light/<br>2 h dark | 20 h light/<br>4 h dark | 16 h light/<br>8 h dark | 12 h light/<br>12 h dark | 8 h light/<br>16 h dark |
| --- | --- | --- | --- | --- | --- | --- |
| WT | 0% | 0% | 0% | 0% | 0% | 0% |
| *dldg1-1* | 100% | 92% | 75% | 0% | 0% | 0% |

**Table S2.**
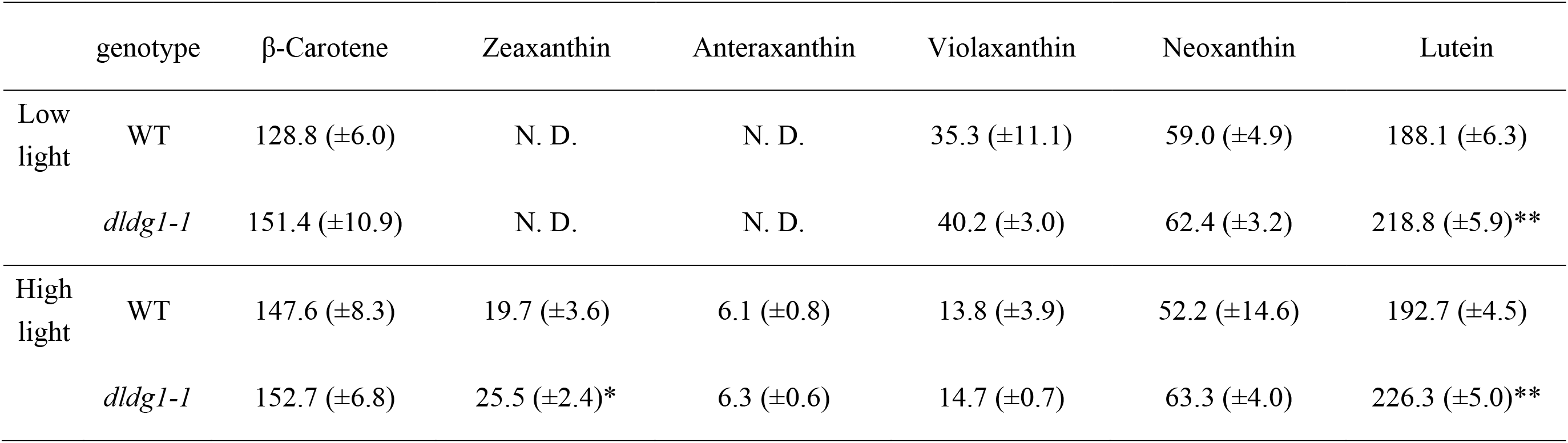
Carotenoid content of WT and *dldg1-1*. For the low-intensity light samples, plants were harvested after exposure to 40 µmol photons m^−2^ s^−1^. For the high-intensity light samples, plants were harvested after exposure to 1,900 µmol photons m^−2^ s^−1^. Carotenoids were detected by HPLC and their amounts normalized to the amount of Chl *a* (mmol/mol Chl *a*). Data are the mean ± S.D. (*n* = 4). \**P* < 0.05, \*\**P* < 0.001 compared with the value for WT under the same light-illumination conditions. N.D., not detected.

**Table S3.**
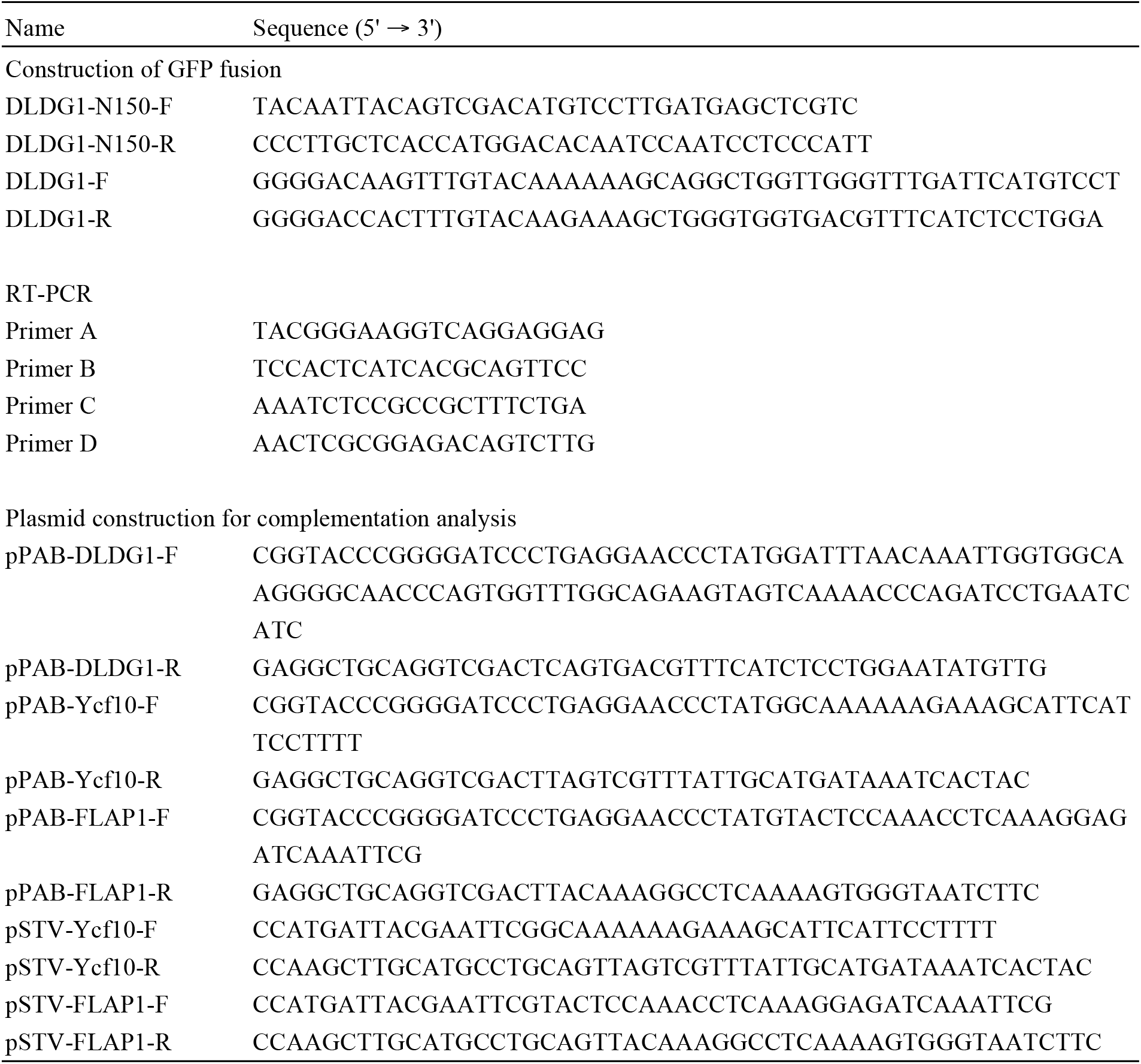
Primers used in this study

